# Hunting with catapults: the predatory strike of the dragonfly larva

**DOI:** 10.1101/2020.05.11.087882

**Authors:** Sebastian Büsse, Alexander Koehnsen, Hamed Rajabi, Stanislav N. Gorb

## Abstract

Dragonfly larvae capture their prey with a strongly modified -extensible- mouthpart using a biomechanically unique but not yet understood mechanism. The current opinion of hydraulic pressure being the driving force of the predatory strike can be refuted by our manipulation experiments and reinterpretation of former studies. On this fact, we present evidence for a synchronized dual-catapult system powered by two spring-loaded catapults. The power output of the system exceeds generally the maximum power achievable by musculature. Energy for the movement is stored by straining a resilin-containing structure at each joint and possibly the surrounding cuticle which is preloaded by muscle contraction. To achieve the precise timing required to catch fast-moving prey, accessory structures are used to lock and actively trigger the system, ensuring the synchronisation of both catapults. As a proof of concept, we developed a bio-inspired robotic arm resembling the morphology and functional principle of the extensible mouthpart. Our study elucidates the predatory strike of dragonfly larvae by proposing a novel mechanism, where two synchronized catapults power the ballistic movement of prey capturing in dragonfly larvae – a so-called synchronized dual-catapult system. Understanding this complex biomechanical system may further our understanding in related fields of bio inspired robotics and biomimetics.

**One Sentence Summary:** The synchronized dual-catapult, a biomechanically novel mechanism for the ballistic movement of prey capturing in dragonfly larvae

## Introduction

Throughout all animal groups, predator–prey relationships often cause an arms race that can lead to the development of elaborate biomechanical mechanisms (*1*). These mechanisms often relay on very fast movements (ballistic movements), for example in prey capturing or jumping (latter most often used as defensive escape mechanism; *2–6*). As there is an inverse relationship between the force output and contraction speed of a muscle, movements relying on high acceleration cannot be driven directly by muscle power. The power output, thus, is modulated to a degree far surpassing the maximal power of a muscle (7–10) due to the instantaneous release of stored energy (*7*,*11*). In many cases these ballistic movements are enabled by a catapult mechanism, where a spring is locked in position and slowly preloaded (for example via muscle contraction). The stored energy is released almost instantaneously via a trigger mechanism (*11*) - the simplest catapult systems is a slingshot. One example of these complex movements is the defensive escape jump of froghoppers (Insecta: Cicadomorpha; *4, 6*). Here a catapult-like elastic mechanism is used to perform one of the fastest jumps known by using chitinised cuticle as a spring (*3*). The elastic protein resilin rapidly returns the leg to its original shape after a jump (using the jumps energy) and allows for repeated jumping (6). Resilin represents an essential element of high resilience, low fatigue, and damping mechanisms in arthropods (*12*) due to its viscoelastic properties (*13*). In the specific case of a catapult system, the near-perfect resilience (92–97%) and a fatigue limit of over 300 million cycles (*14*) in combination with the ability to stretch to over three times its original length and recoil to its initial state without plastic deformation (*15,16*) become important.

Our example here, is the predatory strike Odonata (dragonflies and damselflies) larvae use to capture prey – they evolved a strongly modified, extensible mouthpart called prehensile labial mask (Fig. 1A; *17,18*). These larvae are key predators in their freshwater habitats, hunting invertebrates as well as small vertebrates like tadpoles or fish from an ambush (*19*). These insects can project their specialised mouthpart towards the prey, enabling the larvae to hunt effectively (see supplementary movie S1; *18*). Previous investigations concluded that the protraction of the extensible mouthpart (prehensile labial mask) is partially driven by hydraulic pressure (*20–24*). Strong abdominal dorso-ventral muscles in a rectal chamber – which is also used for respiration – can compress this chamber, thus generating pressure (*25, 26*). By compressing the chamber, a water jet is ejected that propels the larvae forwards (*25, 26*) – this, so-called jet propulsion, represents a special escape behaviour similar to that of squids (*27*) – and was supposed to be redirected and used for the predatory strike (20–24). However, already Tanaka and Hisada suggested a combination of hydraulic pressure and co-contraction of the power muscles as driving force for the predatory strike (*23*). Yet they conducted electrophysiological experiments with no active muscle during the protraction of the mask (see details below). Even more, muscle dissecting experiments (*23*) as well as the presence of specialised morphological structures resembling a locking mechanism (*28*), suggest the necessity of a reinterpretation of the entire system (see also *29*).

**Figure 1:**
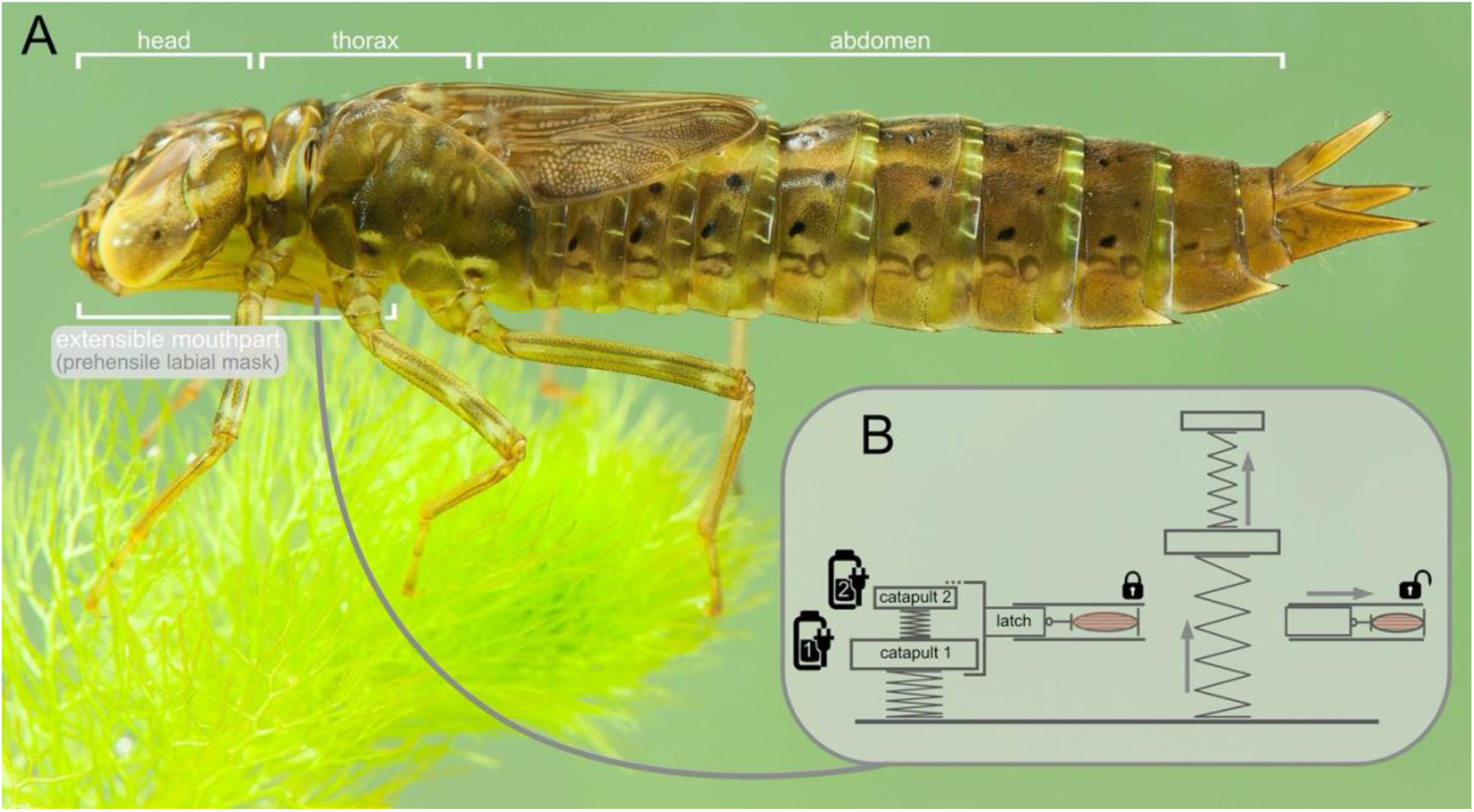
Dragonfly: *Anax imperator* (Odonata: Anisoptera) **A.** Photography, lateral view, © Christophe Brochard – Brochard Photography. **B.** Abstract principle of the concept of a dual-catapult system.

We present evidence for a synchronised dual-catapult system as the driving force of the predatory strike in dragonfly larva. Energy provided by the rather slow contraction of muscles is stored by deformation of two cuticular structures. These two catapults are connected by a joint and operated together (Fig. 1B) to allow for the mentioned power modulation. We could show that the power output required to achieve the observed angular acceleration of the extensible mouthpart (prehensile labial mask) of dragonfly larvae exceeds the power output achievable by the associated musculature (8–10) and therefore making a purely muscle driven movement impossible. Furthermore, our manipulation experiments refute the hypothesis of hydraulic pressure as the driving force (*20–24*). These findings, combined with our morphological data and the bio-inspired robotic arm as proof of concept, provide compelling evidence for the hypothesis, that the extensible mouthpart (prehensile labial mask) is driven by a dual catapult mechanism.

To understand such complex biomechanical system, highlights the evolutionary diversity of insects and often leads to advances in the fields of bio-inspired robotics and biomimetics - as our robotic arm may suggest.

## Results & Discussion

### General morphology and material composition

The extensible mouthpart (prehensile labial mask) of dragonfly larvae in general is a highly modified apomorphic character (*17,18*) – so, a unique character for dragon- and damselflies; which is developed as prey capturing device (*19*). The overall structure consists of a segment 1 (prementum) and 2 (postmentum), which are connected via a cubital-like hinge joint – the connecting joint (prementum-postmentum joint; p-p joint) – allowing uniaxial rotation of both segments relative to each other (Fig. 2A-C, 3; *18*). The prehensile labial mask is connected to the head capsule ventrally via the segment 2 (postmentum) by a membranous joint-like suspension – the head joint (postmentum-head joint; p-h joint) – allowing uniaxial rotation of the extensible mouthpart (prehensile labial mask) relative to the head (Fig. 2A-C, 3; *18*). The connecting joint (p-p joint) consists of large membranous areas, likely supplemented with resilin (*28*). For a detailed description on the morphology and/or material composition of the mouthparts of dragonflies we refer to (*18,28*).

**Figure 2:**
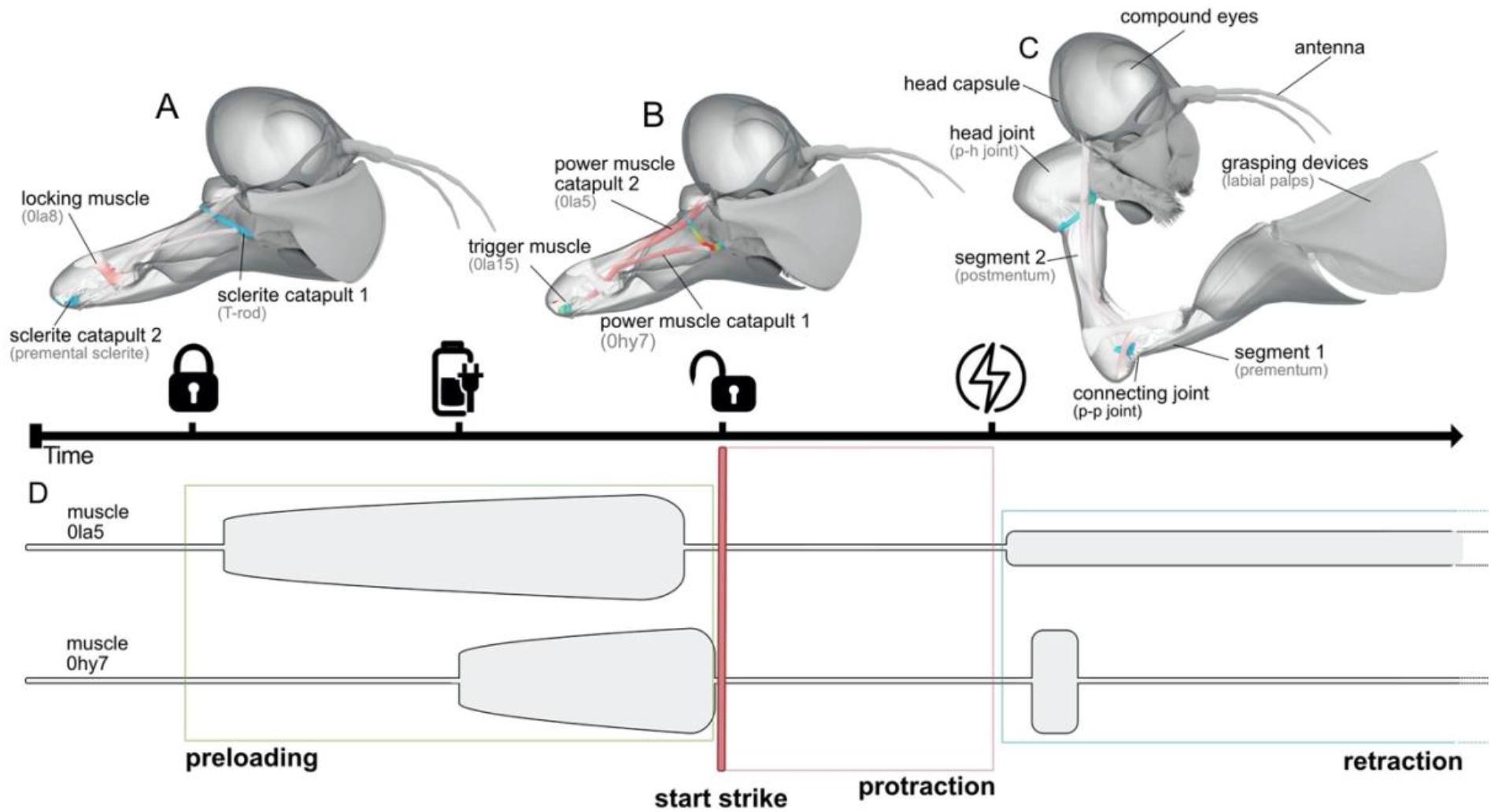
Morphology of the extensible mouthpart (prehensile labial mask) and a simplified sequence of the predatory strike; three-dimensional visualisation derived from μCT data of *Sympetrum* sp. (Odonata: Anisoptera). Colour-code: grey muscles = relaxed, red muscles = contracted; blue sclerites = undeformed, red sclerites = deformed. **A.** preparation for predatory strike, locking **B.** Preloading and triggering of the protraction **C.** Unlocking and protraction of the extensible mouthpart (prehensile labial mask). **D.** Electrophysiology of the predatory strike, showing muscle activity of the power muscles (0la5 and 0hy7) during preloading, protraction and retraction (modified after *23*).

### Introduction of the synchronized dual-catapult system

We propose a novel biomechanical mechanism for the protraction of the extensible mouthpart (prehensile labial mask): a synchronised dual-catapult system, consisting of two catapults activated simultaneously as the main driving force for the predatory strike of dragonfly larvae (Fig. 1B). Both catapults are spring-loaded (*7,11*), generating the main power for the strike by storing elastic strain energy which can be rapidly converted into kinetic energy to enable the high-speed movement. The first catapult provides the power to move the entire extensible mouthpart (prehensile labial mask) towards the prey, with the head joint (p-h joint) as the pivot of rotation (Fig. 2, 3). The second catapult opens the connecting joint (p-p joint; Fig. 2, 3), moving the segment 1 (prementum) downwards and enabling the extensible mouthpart (prehensile labial mask) to unfold (see supplementary movies S1 and S2). In both catapult systems an internal structure is used to store the energy generated by relatively slow action of muscles (Fig. 3, 4A-C). Both structures contain considerable amounts of the viscoelastic protein resilin (*13*) (Fig. 4B, C) which might represent an essential component of this system; however, the surrounding cuticle might play a major role in energy storage as well (cf. *30,31*). To allow energy storage in spring-loaded catapult systems, a lock is needed to prevent untimely release (Fig. 4; *11*). Here, a complex latch mechanism can be found: a clamp, consisting of a groove and a knob that can engage with each other, is locked by a wedge (Fig. 4D-J); locking the system to enable spring loading when needed (Fig. 3). To ensure precise timing of the predatory strike, this dual catapult also needs an active trigger, which is represented by a muscle that can remove the mentioned wedge (Fig. 3C, 4H-J). The two parts of the clamp slide apart and this movement will change the angle of attack (Fig. 4H-J), so that the energy stored in the system will rotate the pivot and cause the catapult arms (segment 1 and 2) to snap forward (Fig. 2,3).

**Figure 3:**
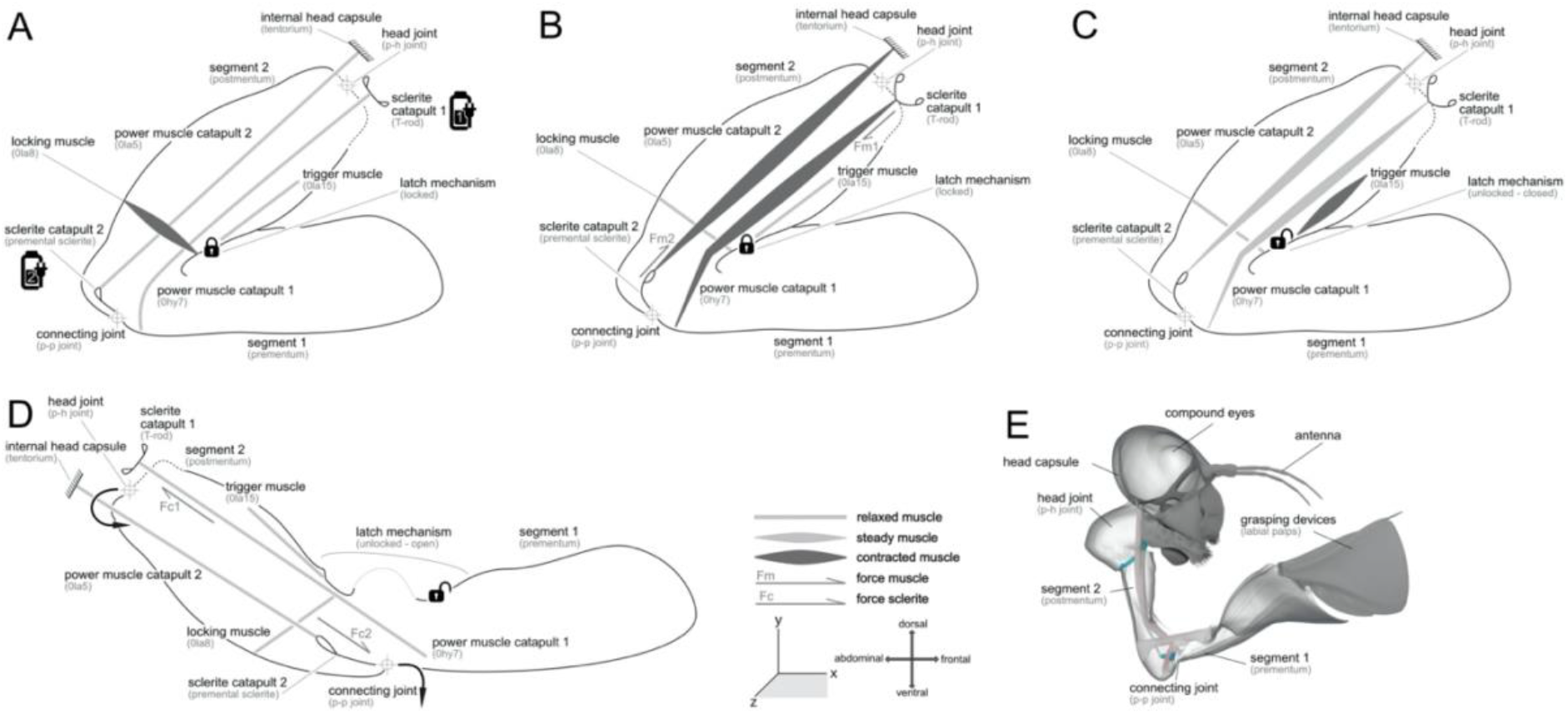
Detailed principle of movement of the extensible mouthpart (prehensile labial mask) and functioning of the synchronised dual catapult system. **A.** Locking: contraction of the locking muscle (0la8) and closing the knob, groove and wedge system (see fig. 4D). **B.** Preloading: contraction of power muscle catapult 1 (0hy7) and deflection (fm1) of sclerite catapult 1 (T-rod) as well as contraction of power muscle catapult 2 (0la5) and deflection (fm2) of sclerite catapult 2 (premental sclerite). **C.** Triggering: contraction of the trigger muscle (0la15) and opening the knob, groove and wedge system (see fig. 4J). **D.** Protraction: releasing the stored energy (fc1 and fc2) of sclerite catapult 1 (T-rod) and sclerite catapult 2 (premental sclerite) to protrect the extensible mouthpart (prehensile labial mask). E. Morphology of the extensible mouthpart (prehensile labial mask), three-dimensional visualisation derived from μCT data of *Sympetrum* sp. (Odonata: Anisoptera).

**Figure 4:**
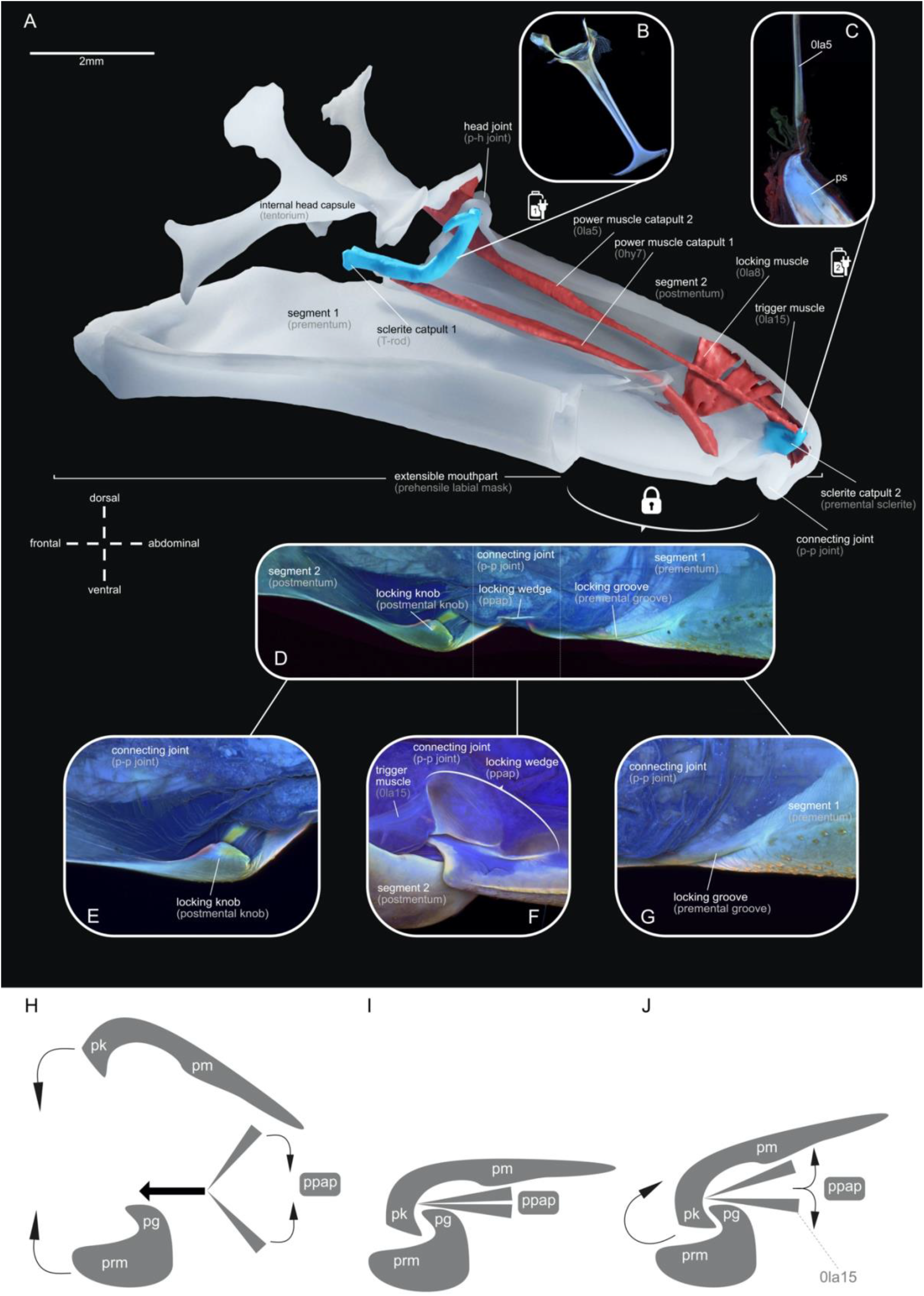
Morphology and material composition of the power unit and locking mechanism of the dual-catapult system in *Anax* sp. (Odonata: Anisoptera). **A.** Extensible mouthpart (prehensile labial mask), three-dimensional visualisation derived from μCT data. **B-G.** CLSM maximum intensity projection, autofluorescences indicates material composition of the cuticle: red – sclerotised, green – chitinous, and blue – resilin-dominated (*28*). **B.** Sclerite catapult 1 (T-rod). **C.** Sclerite catapult 2 (premental sclerite) and power muscle catapult 2 (0la5). **D.** Locking mechanism of the labial mask, dorsal view. **E.** Locking groove (premental groove), detail, dorsal view. **F.** Locking wedge (prementum-postmentum articulatory plate), detail, lateral view. **G.** Locking knob (postmental knob), detail, median view. **H-J.** Principle of the locking/unlocking process. **H.** locking. **I.** locked **J.** unlocking. **Abbreviations**: pg – premental groove (locking groove); pk – postmental knob (loking knob); pm – postmentum (segment 2); ppap – prementum-postmentum articulatory plate (locking wedge); prm – prementum (segment 1).

To allow for a better understanding of this complex biomechanical process, this mechanism with all the structures involved is visually presented and comprehensively explained in a 3D-animation (supplementary movie S2) representing the here proposed biomechanical hypothesis. Furthermore, a detailed description of all the involved structures can be found in the following.

### The dual-catapult system in detail

The dual-catapult system can be subdivided into two interconnected catapults with a single lock (Fig. 3,4), with one mechanism unfolding segment 1 (prementum) and segment 2 (postmentum) - catapult 2 - and one mechanism projecting the entire extensible mouthpart (prehensile labial mask) - catapult 1. At the catapult 1, the sclerite catapult 1 (T-rod) is responsible for storing energy provided by the power muscle of catapult 1 (0hy7; fig 2-4). The spring of catapult 1 (T-rod) is a small sclerite, and its resilin-dominated material composition suggests flexible and resilient properties (Fig. 4 A, B, supplementary figure S4). At catapult 2, energy provided by the power muscle catapult 2 (0la5) is stored in the spring of catapult 2 (premental sclerite; fig 2-4), which is also resilin-dominated (Fig. 4A, C, supplementary figure S4). After the latch system has been locked by the locking muscle (0la8; fig. 3A), the power muscle of catapult 1 (0hy7) deflects the spring of catapult 1 (T-rod; force fm1 in fig. 3B). Simultaneously, the spring of catapult 2 (premental sclerite) is deflected by the power muscle of catapult 2 (0la5; force fm2 in fig. 3B). To simulate the deformation of the sclerites we used a muscle relaxant agent (MgCl_2_) and a muscle contraction agent (KCl). We were able to show that muscle contraction induces a “loaded spring condition” (Supplementary figure 2 D, F) while muscle relaxation induces an “unloaded spring condition” (Supplementary figure 2 E, G) (see also Materials & Methods section). The fact that muscle contraction indeed deforms the sclerites corroborates our hypothesis, that these structures are involved in the described catapult system and are acting as energy storage devices, likely in combination with the surrounding cuticle. This kind of cuticle deformation that is used for energy storage is for example described for trap jaw ants (cf. *31*), here the entire head is deformed by approximately 6% to allow for their powerful mandiular strike.

To lock the extensible mouthpart (prehensile labial mask) during preloading and trigger a strike, we propose an active latch mechanism at the connecting joint (p-p joint). The latch mechanism is composed of: i) the locking groove (premental groove; cf. *28*), which is present on segment 1 (prementum; fig. 4D,G), ii) the locking knob (postmental knob; cf. *28*), which is the counterpart to the groove on the corresponding area of segment 2 (postmentum), – i) and ii) together forming a clamp (Fig. 4D,E), iii) the locking wedge (p-p articulatory plate; cf. *28*) within the connecting joints (p-p joint) articulation (Fig. 4D,F), and iv) the trigger muscle (0la15; fig. 3A). Firstly, the contraction of the locking muscle (0la8; fig. 3A, 4A) provides the energy to actively push the knob over the groove (Fig. 3A,4H), locking the system to enable spring loading when needed. At this point, the locking knob (postmental knob) clamps behind the locking groove (premental groove) and the locking wedge (p-p articulatory plate), is sitting in between, locking these structures like a wedge (Fig. 3A,B, 4I). For a predatory strike, both catapults need to be triggered. Contraction of the trigger muscle (0la15; fig. 3C) triggers the catapults by removing the wedge (p-p articulatory plate) thus forcing the locking groove (premental groove) and the locking knob (postmental knob) to slide apart (Fig. 3C, 4J).

The material composition of these cuticular structures supplement the locking function: the locking wedge (p-p articulatory plate) is divided into two parts with a resilin-dominated ridge at the divide, that allows the folding into a wedge-like sclerite. The locking groove (premental groove) is sclerotised and represents the slot for the locking knob (postmental knob). In turn, the knob serves as the clamp of the latch, it is composed of a sclerotised ridge, at the contact area with the groove; the surrounding resilin-dominated areas allows for movability during locking and unlocking (Fig. 4D-G). In Büsse & Gorb (*28*) the material composition of these parts is described in more detail.

The release of segment 1 (prementum) changes the traction angles of the co-contracting power muscle catapult 1 (0hy7) and power muscle catapult 2 (0la5), causing power muscle catapult 2 (0la5; running above the pivot of rotation of the segment 2 (postmentum)) to lose tension rapidly. Therefore, the power muscle catapult 1 (0hy7; running below the pivot of rotation of the segment 2 (postmentum)), which is connected to the preloaded sclerite catapult 1 (T-rod), is pulling the segment 2 (postmentum) forward. Simultaneously, both the preloaded sclerite catapult 1 (T-rod) as well as the preloaded sclerite catapult 1 (premental sclerite) release the stored modulated power (force fc1 and fc2 in fig. 3D), leading to a projection of the extensible mouthpart (prehensile labial mask).

### Performance of the catapult system

The extensible mouthpart (prehensile labial mask), respectively the two compartments segment 1 (prementum) and segment 2 (postmentum), reach tangential velocities of Ø 0.5 m s-1 and 0.7 m s-1, angular velocities of Ø 71 rad s-1 and 73 rad s-1, tangential acceleration of Ø 40 and 67 m s-2 as well as peak angular acceleration of 5918 and 6674 rad m s-2 (Table 1, supplementary figure S1, S5). For a typical strike, a power output of Ø 2233 W kg-1 and 2114 W kg-1 is necessary to achieve the mentioned performance Here we conservatively calculated the minimum power requirements (we neglect the drag of the system and therefore underestimates the power output); however, the calculated power output surpasses the power output of the fastest-contracting muscles known considerably (8–10). One of the most powerful muscles mentioned in the literature is that of the blue breasted quail (*Coturnix chinensis*) reaching a max. power output during take-off of approximately 400W kg–1 (*32*). The calculated power output for the catapult system powering the predatory strike of dragonfly larvae is intermediate between the lowest power output values for catapult systems using power modulation described, like snow fleas (*33*) with 740 W kg-1 or flea beetles (*34*) with 714 W kg−1 and the most powerful systems like the jumps of froghoppers (*3*) with 3.6×10_4_ W kg−1 or the most powerful predatory strike of mantid shrimps (*5*) with 4.7×10_5_ W kg−1. Confirming that the predatory strike of dragonfly larvae is indeed power modulated.

**Table 1:** Key characteristics of the performance of the predatory strike. Angle: maximum opening angle from resting position for both prementum and postmentum in [°]. ω: maximum angular velocity of both the pre and postmentum during protraction in radians per second. v: maximum tangential velocity at the distal tip of the prementum/postmentum, calculated from angular velocity. α: maximum angular acceleration calculated as first derivative from angular velocity in [rad/s2]. aT: tangential acceleration at the distal tip of prementum/postmentum, calculated from angular acceleration in [m/s^2] and [g]. N = number of biological replicates.

|  | Segment 1 (prementum) |  | Segment 2 (postmentum) |  |  |
| --- | --- | --- | --- | --- | --- |
|  | Mean | SD | Mean | SD | N |
| Angle [°] | 79 | 11 | 126 | 5 | 3 |
| $\omega$ [rad/s] | 72,54 | 33,32 | 71,42 | 26,89 | 5 |
| $v$ [m/s] | 0,73 | 0,33 | 0,49 | 0,18 | 5 |
| $\alpha$ [rad/s<sup>2</sup>] | 6673,50 | 3697,30 | 5917,69 | 3046,10 | 5 |
| $a_T$ [m/s<sup>2</sup>] | 66,74 | 36,97 | 40,24 | 20,71 | 5 |
| $a_T$ [g] | 6,80 | 3,77 | 4,10 | 2,11 | 5 |
| P[W/kg] | 2113,53 | 2535,46 | 2232,81 | 2560,41 | 5 |

### Manipulation experiments and support of the hypothesis

As mentioned before, the power output of the system surpasses the maximum power of a muscle, as already suggested by (*23*). However, previous investigations indicated that the driving power for the protraction of the extensible mouthpart (prehensile labial mask) is based on hydraulic pressure (*20–24*). In our high-speed video experiments, we could show that *Anax* larvae (n = 5) eject a water jet from the rectal chamber (jet propulsion) during the prey capturing process (see supplementary movie S3 part A). The simultaneity of predatory strike and jet propulsion is most likely a mechanism to counter the recoil which originates from the antagonistic force of quickly accelerating the rather large extensible mouthpart (prehensile labial mask; *35*). This observation is further supported by high-speed video recordings of *Sympetrum* larvae, where the larvae show no jet propulsion but a distinct recoil during the prey capturing process (see supplementary movie S3 part B). Larvae of this taxon are partially burrowed in the soil during hunting (*19*) so jet propulsion seems not to be used/needed for recoil prevention. Furthermore, our observations show dragonflies using jet propulsion for moving towards the prey and performing a predatory strike almost simultaneously. These observations are supported by similar findings for other anisopteran species (*36*). On the one hand, the same mechanism cannot be used for both jet propulsion and propelling the prehensile mask – especially because the predatory strike needs a closed abdomen (anal valve) and jet propulsion an open one (*20*). On the other hand, the simultaneity of these processes might explain the peaks in the hydraulic measurements during the prey capturing process in earlier investigations (*21–24*) and therefore the associated misinterpretation that hydraulic pressure is involved in the protraction of the extensible mouthpart (prehensile labial mask).

Furthermore, the study of Tanaka and Hisada (*23*), especially the included electrophysiology, showed impressively that the only muscles capable of moving segment 1 (prementum) and 2 (postmentum) – the power muscle catapult 2 (0la5; extensor - *23*) and power muscle catapult 1 (0hy7; flexor - *23*) – are not active during the protraction of the extensible mouthpart (prehensile labial mask; fig. 2D). Their experiments highly support our findings: i) both power muscles are active before the starting point of the predatory strike, ii) both muscles are inactive during the main power output of the system, the protraction, iii) are active again during the retraction of the extensible mouthpart (prehensile labial mask; fig. 2D; *23*). Even more their muscle dissection experiments (*23*) clearly showed the importance of the extensible mouthparts (prehensile labial mask) musculature and the insignificance of hydraulic pressure. Dissection of either power muscle catapult 1 (0hy7) or power muscle catapult 2 (0la5) causes abnormal strike movements. Especially after dissection of power muscle catapult 1 (0hy7), the head joint (p-h joint) remains immobile, whereas the connecting joint (p-p joint) opens rapidly and the extensible mouthparts (prehensile labial mask) hits the ground (*23*). In this case, the power muscle catapult 1 (0hy7) is not able to preload the spring of catapult 1 (T-rod), thus causing the abnormal strike behaviour – however, this is not affecting catapult 2. Additionally, we performed manipulation experiments, using the physiological effect of MgCl_2_ as muscle relaxant agent (*37*) to manipulate the abdominal muscles of the rectal chamber (*25, 26*) in such a way as to prevent the generation of hydraulic pressure either for jet propulsion or for the protraction of the extensible mouthparts (prehensile labial mask; *20–24*). After injecting MgCl_2_ into parts of the abdominal muscles related to the rectal chamber, *Anax* larvae (n = 5) were able to perform a predatory strike but could not use jet propulsion as an escape mechanism in response to an external stimulus (see supplementary movie S4). As a control, unmanipulated *Anax* larvae (n = 5) showed jet propulsion in direct response to an external stimulus (see supplementary movie S4). This manipulation experiments clearly refuted the hydraulic hypothesis (*21–24*).

The electrophysiology and muscle dissection experiments (*23*) as well as the manipulation experiments completely refuted the hydraulic hypothesis (*20–24*). Also, the combinational hypothesis of Tanaka and Hisada (*23*) where hydraulic pressure and muscular co-contraction is producing the main power for the predatory strike can be refuted. The power output calculations clearly show that muscle contraction alone cannot provide the power necessary for the observed movement. Hence these findings strongly corroborate our hypothesis that the driving power for the protraction of the extensible mouthparts (prehensile labial mask) is generated by a synchronised dual-catapult system.

### Proof of concept

To test whether the hypothesized interplay of muscles, springs and locks can actually generate a predatory strike like motion, we used our detailed morphological findings to create a robotic model of theprehensile labial mask (fig. 5). The μCT-data was used to ascertain general proportions and match the axes of rotation of both head joint (p-h joint) and connecting joint (p-p joint) as exactly as possible. Muscle movement was imitated by servo motors with matched traction angles. The energy storing sclerites were imitated by steel tension springs (see material & methods). Using this setup, based on the described morphology and hypothesized mechanical configuration, we could show that the artificial extensible mouthparts (prehensile labial mask) is moving in a comparable way as the real predatory strike of a dragonfly larvae (see supplementary movie S5). This proof of concept is intriguingly underlining our hypothesis.

**Figure 5:**
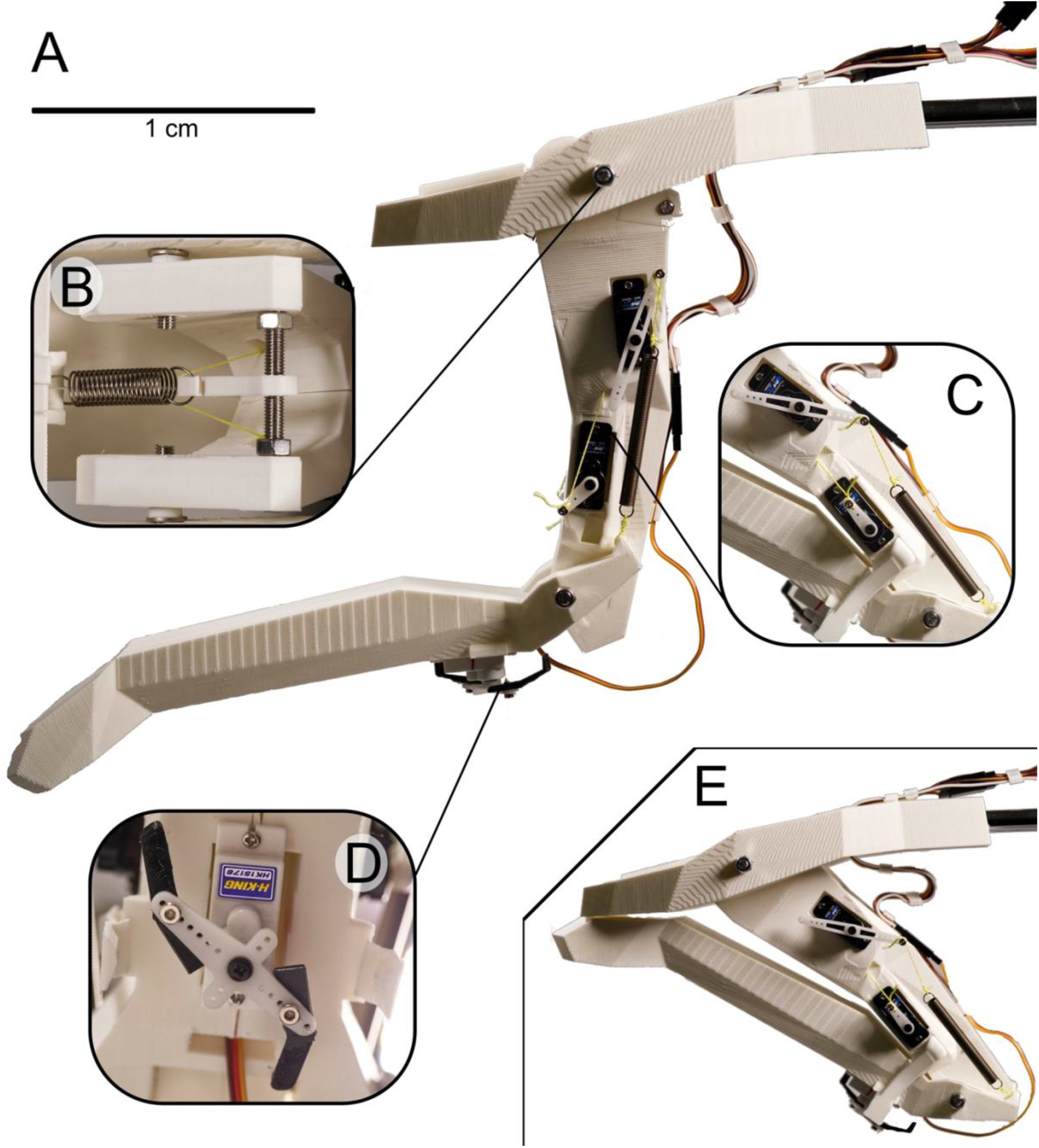
Robotic extensible mouthpart (prehensile labial mask), 3D printed. **A.** open. **B.** artificial spring catapult 1. **C.** artificial spring catapult 2 and artificial muscles (servo motors). D. artificial lock (servo motor). E. closed.

## Conclusion

All in all, we could show that 1) the power output of the system exceeds the maximum power of a muscle and therefore making a purely muscle driven movement impossible; 2) the proposed hydraulic hypothesis as driving force for the predatory strike was a misinterpretation and can be refuted; 3) the present morphology of the extensible mouthparts (prehensile labial mask) represents two catapult systems. The question whether the energy for this high-speed movement is solely stored in the described resilin-dominated sclerites, or parts of the surrounding cuticle are involved as well; requires further research. However, this does not change the functional principle we described here. Our proof of concept using a robotic arm shows impressively the functionality of the proposed mechanism. Our study thus elucidates the predatory strike of dragonfly larvae by proposing a dual-catapult mechanism for the first time. Also highlighting the role of the cuticle as a complex composite material enabling structural integrity and energy storage as one of the main components required for these movements. By implementing two catapults that are triggered together but can be differentially preloaded, this mechanism is probably capable of higher targeting accuracy than other catapults. This makes the prehensile labial mask an intriguing model for further research concerning catapults in biology as well as robotics.

## Materials & Methods

### Animals

Specimens of *Anax* sp. (Anisoptera: Aeshnidae)*, Sympetrum* sp. (Anisoptera: Libellulidae) were collected in Kiel (Germany) in 2016 and 2017 with permission of the ‘Landesamt für Landwirtschaft, Umwelt und ländliche Räume Schleswig-Holstein’.

### Micro-computed tomography

Prior to micro-computed tomography (μCT) analysis, specimens (n=3 per treatment) were fixed for high tissue preservation in alcoholic Bouin solution (= Duboscq-Brasil). We used 3 different treatments prior the tissue preservation: i) for muscle relaxion, pre-fixation in magnesium chloride (MgCl_2_), ii) for muscle contraction, pre-fixation in potassium chloride (KCl) and iii) no pre-fixation. To guarantee that the fixation with KCL as well as MgCl_2_ is not causing artefacts we incubated a test samples (n=3 per structure) for 48h and measured their dimensions using an optical-three-dimensional measuring microscope (VR-3000 series, Keyence, Osaka, Japan; supplementary figure S3). Prior to scanning the samples were dehydrated in an ascending ethanol series and critical-point dried (Quorum E3000; Quorum Tech Ltd., Laughton, UK). For μCT, the critical-point dried samples were mounted on a device-specific specimen holder and scanned (SkyScan 1172; Bruker micro-CT, Kontich, Belgium) with high-resolution settings (40 kV, 250 μA and 0.25° rotation steps, performing a 360° scan). Segmentation and processing of the μCT-data were carried out with Amira 6.0.1 (FEI SAS, Lyon, France). The segmented data were exported as Wavefront ‘.obj’ files for further processing. For three-dimensional visualisation, textures and material shaders for rendering were applied using the open source 3D creation suite Blender (Blender Foundation, Amsterdam, Netherlands, www.blender.org). To Visualise our hypothesis of the predatory strike an armature rig was applied to a CT Data based, retoplogised 3D model and keyframe animation was performed using the high-speed videos as references for correct positioning, angles and timing of the animation. Clips were created using the integrated ‘Cycles’ rendering engine with a resolution of 1920 x 1080p at 25 fps. Animation sequences were saved as ‘Cineon’ image stacks and the final clip was edited in Adobe Premiere Pro CS6 (Adobe Systems Software, San José, CA, USA). Additional 2D animations were created in Adobe After Effects CS6 (Adobe Systems Software, San José, CA, USA).

### Confocal laser-scanning microscopy

All specimens used for confocal laser-scanning microscopy (CLSM) were freshly frozen and stored at −70 °C. The samples were washed in ethanol (100%) and dirt particles were removed using ultrasonic cleaning (Sonorex RK52; Bandelin, Berlin, Germany). The dissected parts were embedded in glycerine (99,99%) on a glass slide and covered with a high-precision cover slip (Carl Zeiss Microscopy, Jena, Germany) prior to scanning. For visualisation, a Zeiss LSM 700 (Carl Zeiss Microscopy, Jena, Germany) was used with the wavelengths 405, 488, 555 and 639 nm and the emission filters BP420–480, LP490, LP560, and LP640 nm. Maximum intensity projections were created using ZEN 2008 software (www.zeiss.de/mikroskopie). For more information on using CLSM to determine the material properties of the insect cuticle, refer to Michels & Gorb (*36*) or Büsse & Gorb (*28*). All images obtained via μCT and CLSM were subsequently processed and combined into figure plates using Affinity Photo and Affinity Design (www.affinity.serif.com).

### Toluidine blue staining

As secondary resilin verification we used a toluidine blue staining (*13,38–41*). The structures (T-rod and premental sclerite) were incubated with 0.1–0.5% toluidine blue (in an aqueous solution of 1% sodium tetraborate) for 30–60 s and destained using glycerin for 48h (see supplementary figure S4). Subsequently analyzed using an optical-three-dimensional measuring microscope (VR-3000 series, Keyence, Osaka, Japan), to detect the bluish stain of resilin containing structures.

### High-speed video recordings

For the high-speed video recordings of the prey capturing process, we used a Photron Fastcam SA1.1 (model 675K-M1; Photron, Pfullingen, Germany) equipped with a 105 mm/1:2.8 macro lens (Sigma, Tokyo, Japan) mounted on a Manfrotto-055 tripod with a Manfrotto-410 geared head (Manfrotto, Spa, Italy) and two Dedocol COOLT3 light sources (Dedotech, Berikon, Switzerland). Settings: 5400 frames per second, exposure time: 1/frame, trigger mode: end, resolution: 1024 × 1024 pixel. Footage was saved as 16-bit TIFF image stacks. Predatory strikes of 5 specimens of *Anax* sp. as well as *Sympetrum* sp. were recorded, with two strikes per individual. Chironomid larvae were manually presented as prey items.

### Motion tracking

Frame by frame data on the position of the prehensile labial mask was obtained from five individuals of *Anax* sp. using the workflow described by Koehnsen et al. (42) From tracking coordinates, the angle between head capsule and postmentum, as well as prementum and postmentum were calculated for every frame. Angular velocity was calculated at every 4th frame and Data was smoothed using an 11th order polynomial (Polynomial regression using R, see also supplementary figure S1b). Angular acceleration was calculated as first order derivative of the obtained curve. Peak velocity and acceleration were obtained from local maxima of the respective curves. Tangential velocity/acceleration at the tip of the prementum/postmentum were calculated from angular velocity/acceleration with the radius r being the average distance from the pivot point to the tip of the respective structure based on all study animals used (n=5). All calculations were done using the open source statistical computation software R Studio (Version 3.3.1 The R Foundation for Statistical Computing, Vienna, Austria).

### Terminology

We will use the term dragonfly(ies) for Odonata (dragonflies + damselflies) for the sake of simplicity. Morphological terminology was used after Büsse et al. (18) and Büsse & Gorb (28). Further, we decide to use the term power modulation rather than power amplification. The latter is widespread within the biomechanics literature, yet it is misleading. The total energy of a system is conserved over time (first law of thermodynamics). The power (and concordantly energy) output of a closed system can therefore not be amplified by means within the system, which the term power amplification suggests. Instead the power output is modulated (43). Energy is stored and later released, leading to an increased peak power output. For more information on the topic we suggest Haldane et al. (42).

### Power output calculations

The prehensile labial mask of *A. imperator*, was modelled as a planar linkage mechanism with two rigid links; postmentum (pm) and prementum (prm). The mechanism was assumed to be pinned at the postmentum-head joint (p-h joint), which works as a fixed rotation axis. The relative rotation of the links was allowed at the prementum-postmentum joint (p-p joint). Supplementary figure 5 represents the system before and during a strike. For the fixed-axis rotation of the system about the postmentum-head joint (p-h joint), we expressed the moment equation as

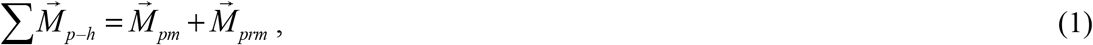

where 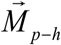 is the total moment needed to accelerate the system about the postmentum-head joint (p-h joint). 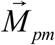 and 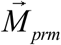 are the moments required to accelerate the postmentum (pm) and prementum (prm) about their joints, respectively.

We first determined 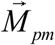 using the following equation

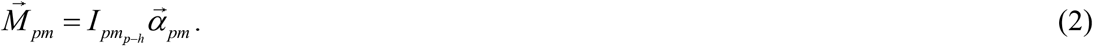

In this equation, 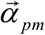 is the vector of the angular acceleration of the postmentum (pm), which was derived from the experimental data. 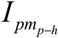 is the mass moment of inertia of the postmentum (pm) about an axis through the postmentum-head joint (p-h joint) and was calculated as:

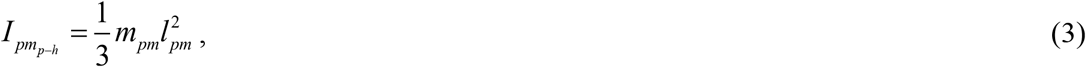

where *m*_*pm*_ and *l*_*pm*_ are the mass and the length of the postmentum (pm).

To calculate 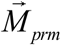, on the other hand, we first determined the linear velocity of the prementum-postmentum joint (p-p joint) using the following equation

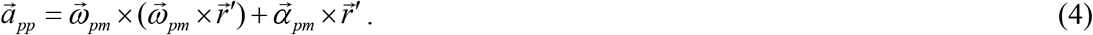

In this equation, 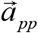 is the vector of the linear velocity of the prementum-postmentum joint (p-p joint). 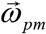 is the vector of the angular velocity of the postmentum (pm) and was measured based on the experimental data. 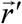 is the position vector of the prementum-postmentum joint (p-p joint).

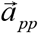, which was calculated using equation (4), was then used to calculate 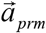, the vector of the linear acceleration of the mass center of the prementum (prm), as follows

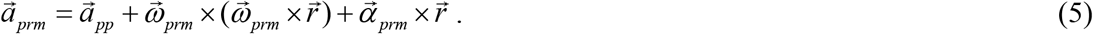

In the above equation, 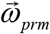 and 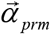 are the vector of the angular velocity of the prementum (prm) and the vector of the angular acceleration of the prementum (prm), respectively. Both 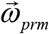 and 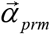 were obtained from experiments. 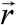 is the position vector of the mass center of the prementum (prm).

After calculating 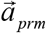 using equation (5), the result was substituted into the moment equation of the prementum (prm) about the prementum-postmentum joint (p-p joint) as follows

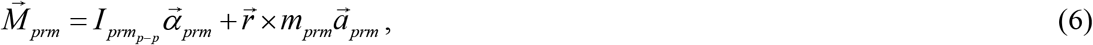

where *m*_*prm*_ is the mass of the prementum (prm) and 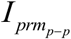 is the mass moment of inertia of the prementum (prm). The latter was determined using the equation below

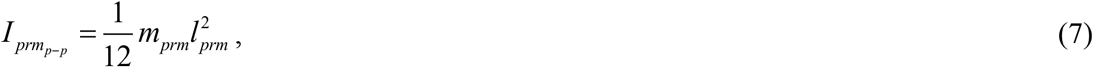

where *l*_*prm*_ is the length of the prementum (prm).

After calculating 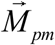 and 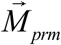 using equations (2) and (6), the results were substituted into the equation below and the power output, *P*, of each of the two muscles (i.e. 0la8 and 0hy7) was determined at different time steps during the strike

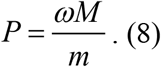

In this equation, *ω* is the magnitude of the vector of the angular velocity of each link, *M* is the magnitude of the moment accelerating the link, and *m* is the mass of the muscle involved in the link rotation. The mass of each muscle was taken from (*23*). The velocity measurements were repeated for the data extracted from five individuals with ten predatory strikes.

### Manipulation experiments

For the manipulation experiments, we injected 2–4 ml of a 20 mmol/l solution of magnesium chloride (MgCl_2_) as muscle relaxant agent (*37*) into the abdominal dorso-ventral muscles of *Anax* sp. larvae (n=5). After 2 to 5 min, the injected dorso-ventral musculature was relaxed and the specimens were not able to produce the necessary hydraulic pressure for jet propulsion (in response to an external stimulus). Chironomid larvae (Insecta: Diptera) were manually presented as prey items. After a successful predatory strike, the larvae were given an external stimulus to trigger an escape reaction. We scored the ability to use jet propulsion as an escape mechanism after a successful predatory strike in manipulated and unmanipulated specimens.

### Artificial prehensile mask (proof of concept)

To proof the concept of a catapult driven prehensile labial mask, that can successfully generate a predatory strike using the herein described morphology, we construct an artificial model. A 3D model was designed using the 3D creation suite Blender (v2.79, Blender Foundation, Netherlands, www.blender.org). Relative proportions, axes of rotation and angles of traction were derived from CT data and hs-video. Individual parts were exported as “.stl” files, and printed on a Prusa i3 Mk2S FDM 3D printer (Prusa Research s.r.o., Prague, Czech Republic) using polylactic acid filament (Prusa Research s.r.o., Prague, Czech Republic). Steel tension springs (One9.1×27.4×1mm spring at the ph-catapult and two 5.7×59.2×.06mm springs at the pp-catapult) serve as energy storage devices. Each catapult is preloaded using two Turnigy MX-M801 servo motors (HexTronics Ltd. Kowloon Bay, Hong Kong). The latch mechanism is triggered using a HobbyKing HK 15178 servo motor (HexTronics Ltd. Kowloon Bay, Hong Kong). Motors are controlled by an Arduino Uno R2 Board (Arduino.cc) using custom code.. The entire system was powered with a 5V 600mA Power Supply Unit (Elegoo Power MB V2, Elegoo Inc, Shenzen, China).

High-speed video footage of the artificial labial mask was captured at 1000fps at a resolution of 1280×1024p using an Olympus i-Speed 3 high-speed camera (iX Cameras, Rochford, Essex, UK) equipped with a Sigma Compact Hyperzoom 28-200mm/1:3.5-5.6 macro lens (Sigma, Tokyo, Japan). Data was saved in the AVI Codec and edited using Adobe Premiere CS6 (Adobe Systems Software, San José, CA, USA).

## Acknowledgments

We are grateful to S. Bodenstein and K. Lehmann for help with the collection of the larvae and to H.-L. Tröger and M. Garitz for undergraduate research assistance. We are especially thankful for the help of L. Heepe for proofreading and many elucidating discussions.

## Author contributions

S.B. and S.N.G. designed the project and developed the concept of the study. A.K. and S.B. did the μCT analysis and post-processing. S.B. and A.K. did the HS-Video recordings. S.B. carried out the CLSM analysis and conducted the manipulation experiments. A.K. did the motion tracking and developed the robotic model. H.R. calculated the power output of the system. All authors wrote the manuscript as well as read and approved the final version.

## Data and materials availability

Supplemental data on reasonable request.

## Supplementary Materials

Fig. S1 to S5 (one .pdf file)

Movies S1 to S5 (on reasonable request).

