## Supplementary Material for "Hunting with catapults: the predatory strike of the dragonfly larva"

bioRχiv
THE PREPRINT SERVER FOR BIOLOGY

**Supplementary Information for:**

### Hunting with catapults: the predatory strike of the dragonfly larva

Sebastian Büsse, Alexander Koehnsen, Hamed Rajabi & Stanislav N. Gorb

Sebastian Büsse


**This PDF file includes:**

Fig. S1 to S5
Legends for Movies S1 to S5

**Other supplementary materials for this manuscript include the following:**

Movies S1 to S5 (on reasonable request)

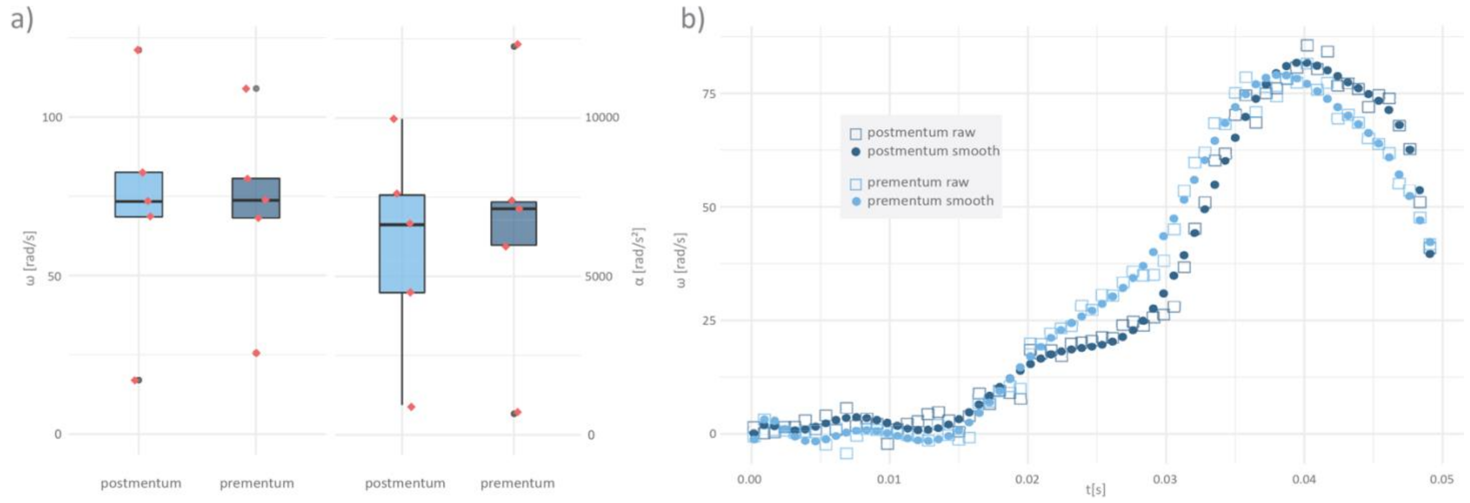


**Fig. S1. Kinematics of the predatory strike based on motion-tracking data.**

**A.** Whisker-plots representing peak angular velocity (left) and peak angular acceleration (right) of the postmentum and prementum respectively (n=5 individuals á 2 strikes). Box ends define 25th and 75th percentiles, with a line highlighting the median and error bars at the 10th and 90th percentiles. Individual data are marked by red dots, outliers are marked by grey dots. **B.** Angular velocity per time of post- and prementum. Squares represent raw tracking-data, dots represent fitted data (11th order polynomial, see material and methods). Exemplary data of one strike.

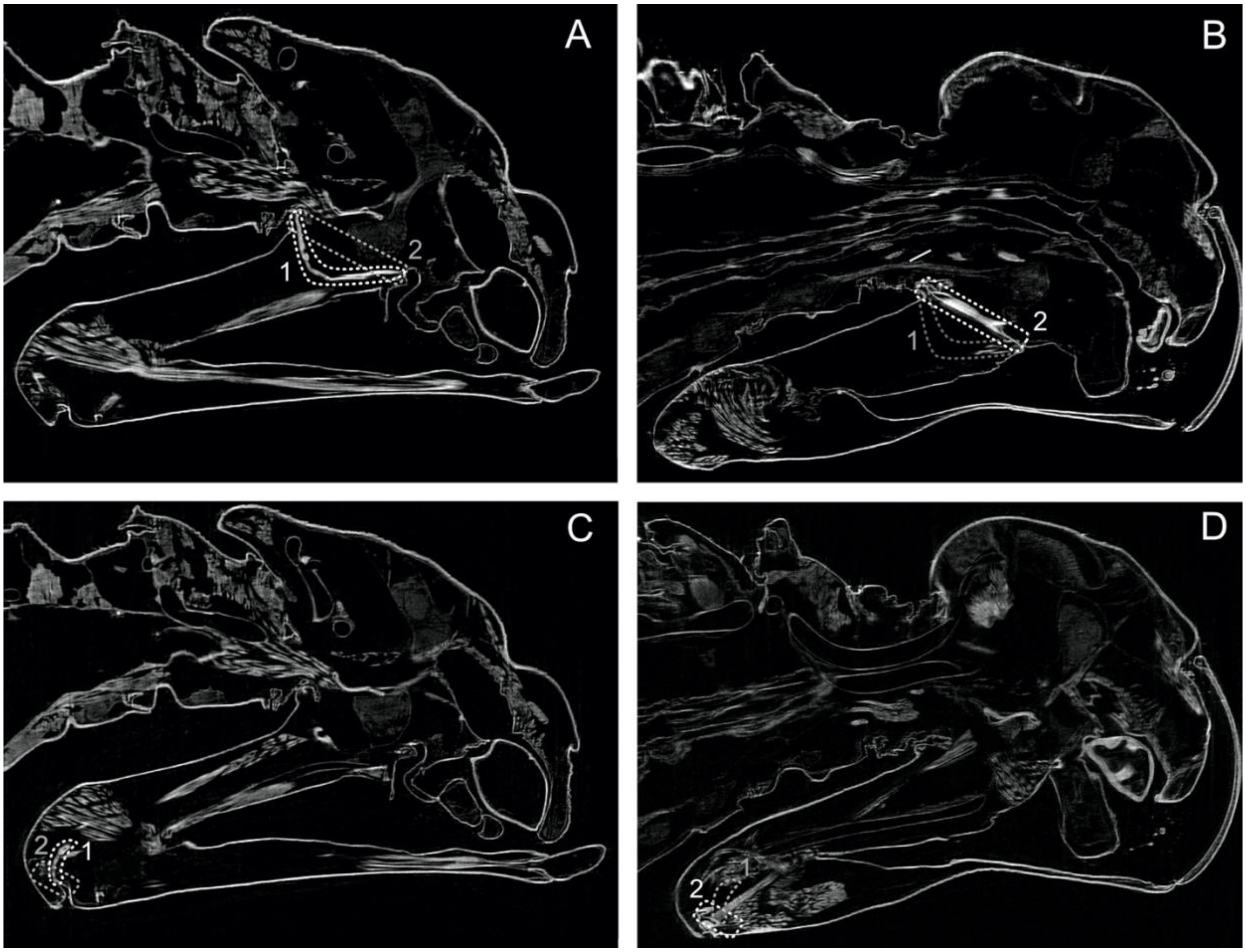


**Fig. S2. Deformation of sclerites using muscle contraction and relaxion agents, virtual slices derived from µCT data, lateral view.**

**A&B.** Focus on the T-rod**. C&D.** Focus on the premental sclerite. **A-D.** In broken lines: **1** showing the area of the bended T-rod respectively the premental sclerite. **2** showing the area of the straight T-rod respectively the premental sclerite. **A&C.** Musculature contracted (KCl) resembling the loaded catapult **B&D** Musculature relaxed ($MgCl_2$) resembling the unloaded catapult.

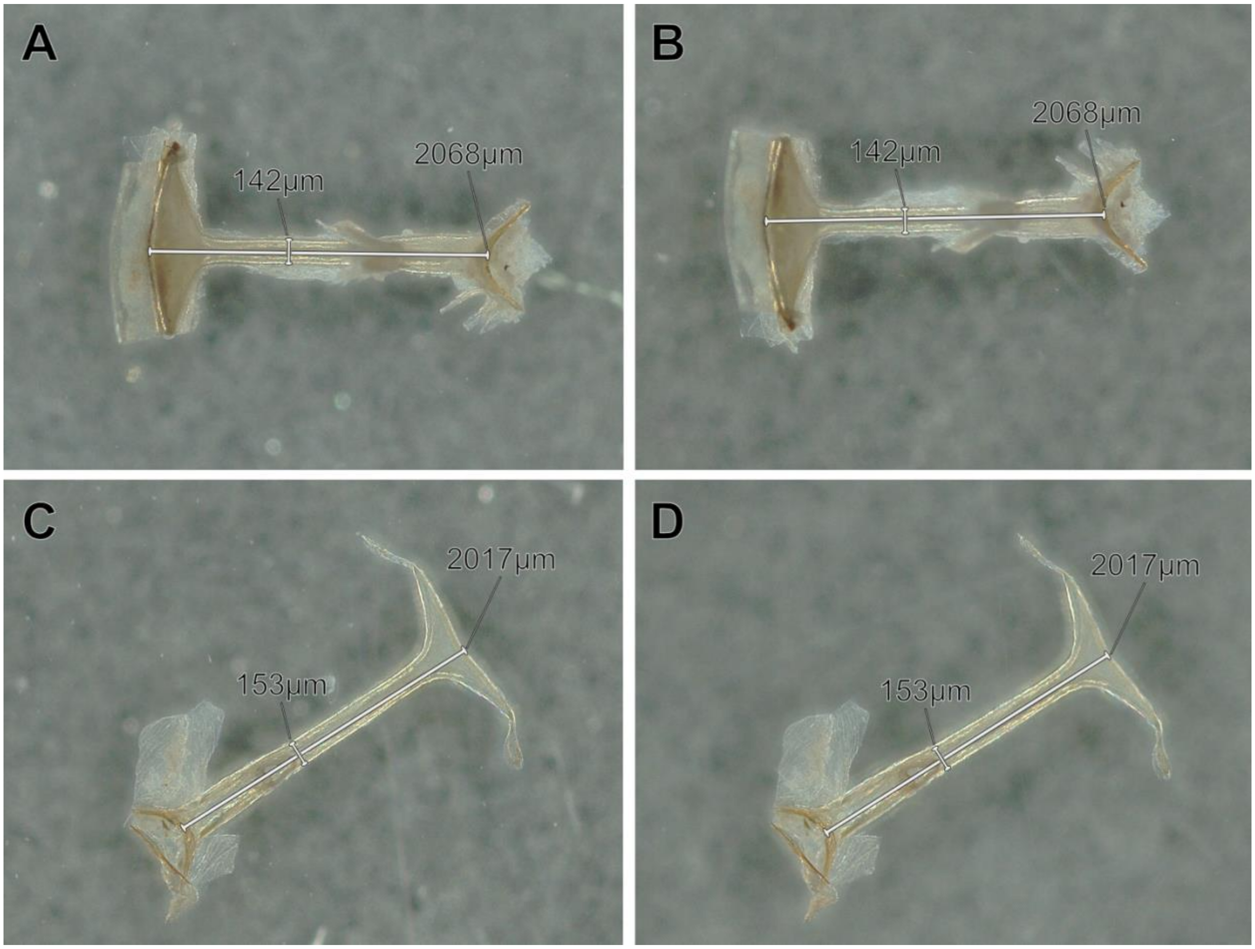


**Fig. S3. Artefacts control of KCL and $MgCL_2$ fixation of the T-rod.**

**A-D.** Exemplary test samples (n=3 per structure), T-rod, measured dimensions using an optical-three-dimensional measuring microscope. **A&C.** freshly dissected. B&D. after 48h incubation **B.** KCL **D.** $MgCl_2$.

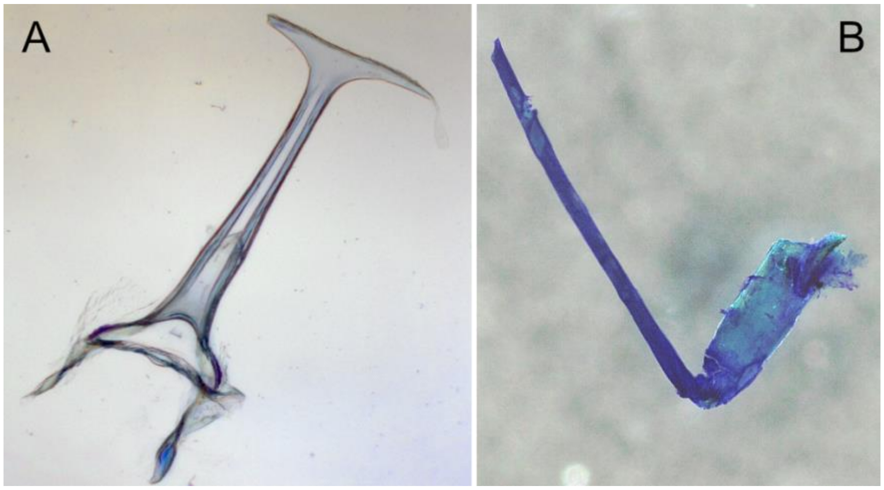


**Fig. S4. Toluidine blue staining as secondary resilin verification.**

**A&B.** Incubated with 0.1–0.5% toluidine blue (in an aqueous solution of 1% sodium tetraborate) for 30–60 s and destained using glycerin for 48h. **A.** T-rod. **B.** Premental sclerite.

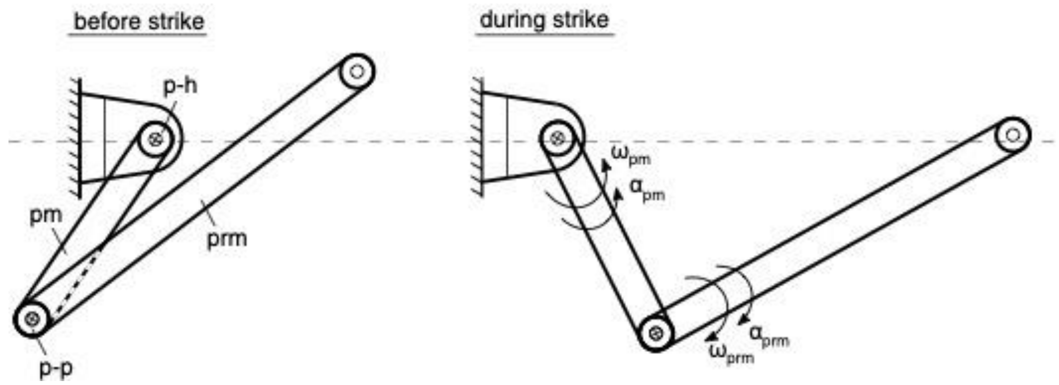


**Fig. S5.** Schemes of the predatory strike of Odonata larvae. Before strike and during the strike.

### Legends for supplementary Movies

**Movie S1.** (separate file)
*Anax* sp. (Odonata: Anisoptera), high-speed videography of the predatory strike, 5400 fps.

**Movie S2.** (separate file)

Fictive 3D animation of the predatory strike, based on CT-data.

**Movie S3.** (separate file)

*Anax* sp. (Odonata: Anisoptera), high-speed videography of recoil prevention, 5400 fps.

**Movie S4.** (separate file)

*Anax* sp. (Odonata: Anisoptera), videography of the manipulation experiments.

**Movie S5.** (separate file)

Artificial prehensile mask, videography of the proof of the concept
